# Metagenomics and quantitative stable isotope probing offer insights into metabolism of polycyclic aromatic hydrocarbons degraders in chronically polluted seawater

**DOI:** 10.1101/2020.10.16.343509

**Authors:** Ella T. Sieradzki, Michael Morando, Jed A. Fuhrman

## Abstract

Bacterial biodegradation is a significant contributor to remineralization of polycyclic aromatic hydrocarbons (PAHs): toxic and recalcitrant components of crude oil as well as byproducts of partial combustion chronically introduced into seawater via atmospheric deposition. The Deepwater Horizon oil spill demonstrated the speed at which a seed PAH-degrading community maintained by low chronic inputs can respond to acute pollution. We investigated the diversity and functional potential of a similar seed community in the Port of Los Angeles, a chronically polluted site, using stable isotope probing with naphthalene, deep-sequenced metagenomes and carbon incorporation rate measurements at the port and in two sites further into the San Pedro Channel. We demonstrate the ability of a local seed community of degraders at the Port of LA to incorporate carbon from naphthalene, leading to a quick shift in the microbial community composition to be dominated by these normally rare taxa. We were able to directly show that assembled genomes belonged to naphthalene degraders by matching their 16S-rRNA gene with experimental stable isotope probing data. Surprisingly, we did not find a full PAH degradation pathway in those genomes and even when combining genes from the entire microbial community. We analyze metabolic pathways identified in 29 genomes whose abundance increased in the presence of naphthalene to generate metagenomic-based recommendations for future optimization of PAHs bioremediation.

**Importance:** Oil spills in the marine environment have a devastating effect on marine life and biogeochemical cycles. Oil-degrading bacteria occur naturally in the ocean, especially where they are supported by chronic inputs of oil, and have a significant role in degradation of oil spills. The most recalcitrant and toxic component of oil is polycyclic aromatic hydrocarbons. Therefore, the bacteria who can break those molecules down are of particular importance. We identified such bacteria at the port of Los Angeles, one of the busiest ports worldwide, and characterized their metabolic capabilities. Based on those analyses we proposed chemical targets to stimulate the activity of these bacteria in case of an oil spill in the port of LA.

## Introduction

Polycyclic aromatic hydrocarbons (PAHs) are recalcitrant, mutagenic and carcinogenic components of crude oil as well as byproducts of incomplete combustion (1). Microbial biodegradation has an important role in PAH remediation alongside physical weathering processes (2). Biodegradation of PAHs captured much scientific attention after the Deepwater Horizon (DWH) oil spill in the Gulf of Mexico in 2010. Several studies measured PAH degradation rates (3, 4) and showed enrichment of known PAH-degrading bacteria in beaches, surface water, the deep-sea plume and sediments even months after the spill began (5–9). Bacteria known to have the ability to utilize PAHs as a carbon source include strains of *Cycloclasticus, Colwellia, Pseudomonas, Alteromonas*, and others (6, 10–12). Many coastal sites worldwide experience chronic input of PAHs, mainly from atmospheric deposition and natural oil seeps. Recent studies show that chronic pollution supports a consistent “seed” of PAH degrading bacteria which can respond quickly to acute pollution such as an oil spill (6, 10–15). Stable isotope probing (SIP) is a well-established method for the identification of environmental bacteria utilizing targeted substrates, in this case PAHs (16, 17), in which ^13^C PAHs are added to samples in order to make the DNA of PAH-utilizers heavy and thus capable of physical separation in a density gradient. A large-scale SIP study was performed on DWH surface and deep-plume water, revealing local strains of PAH-degrading bacteria that responded to the input of hydrocarbons (6). Some genomes of those bacteria were assembled from mesocosm metagenomes in order to further explore their PAH metabolism (18). The studies mentioned above, like many others, focus only on the high-density (i.e. most heavily ^13^C-labeled) fractions under the assumption that the most heavily labeled organisms, and thus the main targets, will be found there. However, this strategy may lead to overlooking degraders with low-GC (i.e. naturally lower DNA density) genomes, whose DNA may not appear in the heaviest fractions even if they include moderate amounts of ^13^C (19). For example, 50% enrichment of a genome with 30% GC using ^13^C would result in a mean weighted density of 1.7175 g*ml^-1^ (20), whereas the center of gradient is commonly set at 1.71-1.72 g*ml^-1^. Tag-SIP is a powerful and particularly sensitive extension of the standard SIP approach, in which DNA from both labeled samples and parallel unlabeled controls are separated into density fractions, and the 16S-rRNA gene is amplified from all density fractions of both samples for comparison. This approach, circumventing GC-based bias, allows us to track substrate incorporation by a single taxon demonstrated by an increase (shift) in its DNA density in the labeled samples compared to controls (21, 22).

One of the main motivations to study PAH-degrading organisms is to characterize their metabolic requirements. Understanding the suite of nutrients and cofactors those organisms require could potentially be applied towards bioremediation and biostimulation (23). The combination of SIP with metagenomics can help reveal metabolic dependencies within assembled genomes of PAH-degraders (2, 15, 18, 23). Additionally, it has been proposed that PAH biodegradation is a community process rather than fully performed by a single taxon (18). Thus, it is important to investigate potential degradation using Tag-SIP and deep sequencing so as to identify not only the function of the organisms that initiate the degradation but also of the other organisms involved in later steps of degradation.

While the Port of LA (POLA) is not routinely monitored for PAHs, it is located in an area with natural oil seeps, it houses a ship-refueling station of high-aromatic-content marine diesel (ETC Canada, 11/22/2017) and it is surrounded by the second largest metropolitan area in the USA, which is likely a source of consistent atmospheric PAH deposition. The LA-Long Beach port is the busiest port in the United States and the 10^th^ busiest in the world according to the international association of ports and harbors (IAPH, 11/26/2017). A report published in 2010 revealed detectable levels of many PAHs at various stations within the port and as far as outside the port entrance, corresponding to our sampling site (24, 25). Due to the high marine traffic at POLA, there is constant resuspension into the water column of sediment, which would normally (without such regular resuspension) serve as a sink for PAHs due to their tendency to attach to particles (24, 25).

Here we provide evidence for the existence of a rare seed PAH-degrading, free-living (0.2-1 μ) microbial community in the chronically polluted Port of Los Angeles. We show that this seed can “spring into action” as a response to high PAH input, and start a degradation process leading to a specialized community dominated by two genera. We then discuss metabolic characteristics of the local PAH-degraders using largely complete genomes of PAH degraders and propose targets for biostimulation experiments in this system.

## Materials and Methods

### Sample collection

Surface seawater was collected in July and October 2014 from three sites across the San Pedro Channel near Los Angeles, CA, USA (fig. 1): the port of Two Harbors, Santa Catalina Island (CAT), the San Pedro Ocean Time-series (SPOT) and the Port of Los Angeles (POLA). An additional sample was collected in May 2015 only from POLA. Water was collected by HDPE bucket into two 10 liter LDPE cubitainers and stored in a cooler in the dark until arrival to the lab.

**Figure 1:**
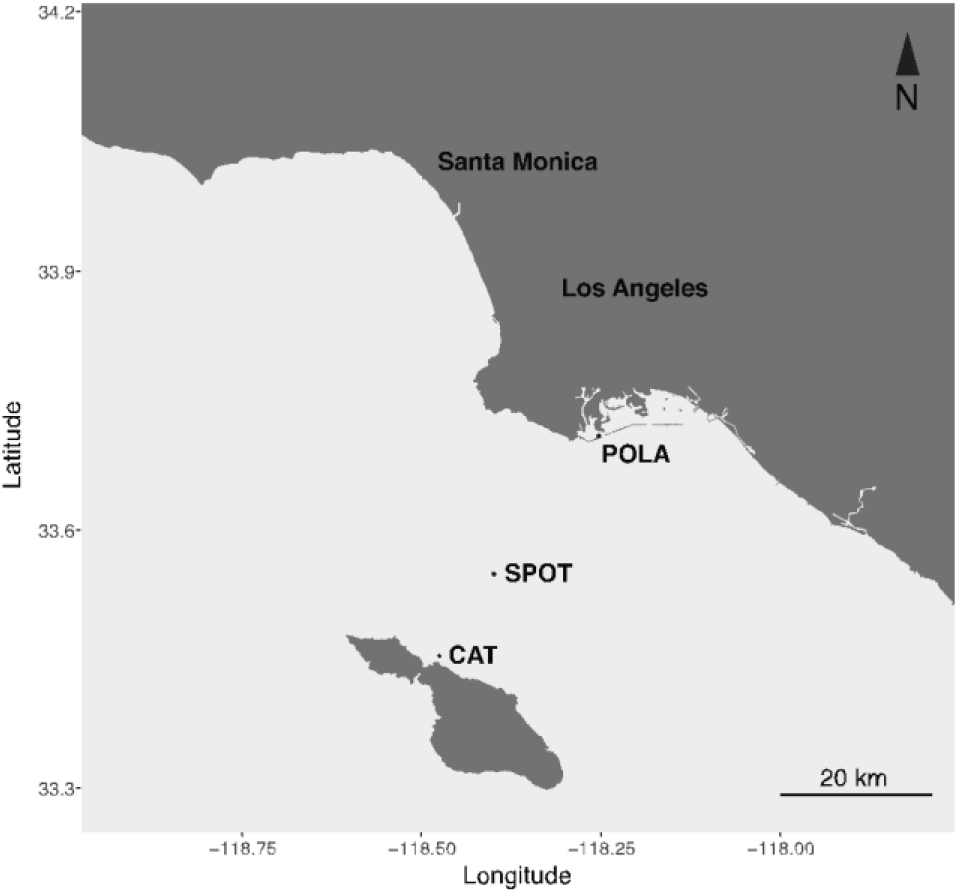
map of the sampling sites across the San Pedro Channel. The sites are within a range of 40 km.

### Isotope addition and incubation

400 nM of either unlabeled (^12^C) naphthalene or fully labeled ^13^C-10-naphthalene (ISOTEC, Miamisburg, OH, USA) were added to each 10 L cubitainer of water collected without replication. This concentration is roughly 3 orders of magnitude lower than the solubility of naphthalene in water (30 mg/L = 234μM). 1μM ammonium-chloride was also added to each bottle to prevent nitrogen limitation of PAH degraders. Naphthalene-enriched seawater from July and October 2014 was incubated in 10% ambient light, which is comparable to the light at 5m depth at SPOT, at surface water temperature (17°C) for 24 hours, and from May 2015 in the dark at surface water temperature (20°C) for 88 hours. Incubation of POLA water in the dark was performed to simulate more closely conditions in that site, as light attenuation at POLA is much steeper compared to the other sites (sup. fig. S1). At the end of the incubation period the seawater was filtered through an 80 nM mesh and a glass fiber Acrodisc (Millipore-Sigma, St. Louis, MO, USA) prefilter (pore size 1 μm) followed by a 0.2μm polyethersulfone (PES) Sterivex filter (Millipore-Sigma) to capture only the free-living microbes. After filtration, 1.5ml Sodium-Chloride-Tris-EDTA (STE; 10mM Tris-HCl, 1mM EDTA and 100mM NaCl) buffer was injected into the Sterivex casing and the filters were promptly sealed and stored in −80°C.

### Carbon incorporation rate measurement

Seawater from all three sites was incubated in quadruplicate 2-liter polycarbonate bottles with 400nM ^13^C labeled naphthalene. 1μM ammonium-chloride was also added to each bottle to prevent nitrogen limitation of PAH degraders. A single bottle was filtered immediately after amendment addition to establish a t_0_ atom% ^13^C of the particulate carbon at each site. The remaining triplicates were incubated in a temperature-controlled room (see above). Incubations were carried out for ~24 h and were terminated by filtration onto pre-combusted (~5 h at 400°C) 47mm GF/F filters (Whatman, Maidstone, United Kingdom). The filters were then dried at 60°C and kept in the dark until analysis. Isotopic enrichment was measured on an IsoPrime continuous flow isotope ratio mass spectrometer (CF-IRMS). IRMS data were corrected for both size effect and drift before being calculated as previously described (26).

### DNA extraction

Total DNA was extracted from the Sterivex filters using a modified DNeasy Plant kit protocol (Qiagen, Hilden, Germany). The Sterivex filters containing STE buffer were thawed. 100 μL 0.1mm glass beads were added into the filter casing and put through two 10-minutes cycles of bead beating in a Tissuelyser (Qiagen, Hilden, Germany). The flow-through, pushed out using a syringe, was incubated for 30 minutes at 37°C with 2 mg/ml lysozyme followed by another 30 minutes incubation at 55°C with 1 mg/ml proteinase K and 1% SDS. The resulting lysate was loaded onto the DNeasy columns followed by the protocol as described in the kit instructions. Only samples from POLA 5/2015 yielded enough DNA for ultracentrifugation of both the labeled and unlabeled DNA. However, metagenomes were sequenced from all dates and sites except SPOT 10/2014.

### Ultracentrifugation and density-fractions retrieval

Isopycnic ultracentrfugation and gradient fractionation were performed as described in previous work (22, 27). Briefly, DNA from the labeled and unlabeled samples was added into separate quick seal 5ml tubes (Beckman Coulter, Indianapolis, IN, USA) combined with 1.88 g/ml CsCl and gradient buffer for a final buoyant density of 1.725 g/ml. The tubes were sealed and centrifuged in a Beckman Optima L100 XP ultracentrifuge and near-vertical rotor NVT 65.2 at 44100 rpm (190,950 rcf) at 20°C for 64 hours.

The gradient was divided into 50 fractions of 100μl each. Refraction was measured using 10 μl of each fraction using a Reichert AR200 digital refractometer, and converted into buoyant density (ρ=10.927*n_c_-13.593) (28). DNA in each fraction was preserved with 200 μl PEG and 1 μl glycogen, precipitated with ethanol, eluted in 50 μl Tris-EDTA (TE) buffer and quantified using PicoGreen (Invitrogen, Carlsbad, CA, USA).

### Amplification of the 16S-rRNA V4-V5 hypervariable regions

PCR was performed on each fraction with detectable DNA. Each reaction tube contained 12 μl 5Prime Hot Master Mix (Quantabio, Beverly, MA, USA), 1 μl (10pg) barcoded 515F-Y forward primer (5’-GTG**Y**CAGCMGCCGCGGTAA), 1 μl (10pg) indexed 926R reverse primer (5-CCGYCAATTYMTTTRAGTTT), 1 ng of DNA and 10μl molecular-grade water. Thermocycling conditions:

1. 3 min denaturation at 95°C
2. 30 cycles of denaturation at 95°C for 45 sec, annealing at 50°C for 45 sec, and elongation at 68°C for 90 sec
3. final elongation at 68°C for 5 min

PCR products from each fraction were cleaned using 1x Agencourt AMPure XP beads (Beckman Coulter), quantified by PicoGreen and diluted to 1 ng/μl. A pool of 1 ng of each uniquely-barcoded product was cleaned and concentrated again with 0.8x Agencourt AMPure XP beads. The pooled amplicons were sequenced on Illumina MiSeq (UC Davis, USA) for 600 cycles. Each pool was also spiked with 1 ng of an even and a staggered mock community in order to assess the sequencing run quality (29). All expected OTUs were found in the observed mock communities and accounted in total for 99.5% of the reads, indicating that there was no unexpected bias in the sequencing run (30).

### Amplicon data processing

The raw reads were quality-trimmed using Trimmomatic (31) version 0.33 with parameters set to LEADING:20 TRAILING:20 SLIDINGWINDOW:15:25 MINLEN:200 and merged with Usearch version 7 (32) with a limit of maximum 3 differences in the overlapping region. The resulting merged-reads were analyzed in Mothur (33) and clustered at 99% identity following the MiSeq SOP (March 20^th^, 2016) (34) with one exception: we found that degapping the aligned sequences, aligning them again and dereplicating again fixed an artifact in which a few abundant OTUs were split due to the alignment, despite 100% identity, due to an additional terminal base between the OTU representative sequences. OTUs with a total of less than 10 reads over all fractions were removed from the analysis. The remaining 2366 OTUs were assigned taxonomy using the arb-silva SINA search and classify tool version 1.3.2 (35).

### Detection of OTU enrichment due to substrate incorporation

Plots of normalized abundance as a function of density were generated in R (https://www.r-project.org/) for the top 200 most abundant OTUs in each sample. The abundance of each OTU was normalized to a sum of 1 across all fractions (eq. 1).

Equation 1: Normalization of OTU abundance per fraction where i=fraction number, Ri=relative abundance of the OTU in fraction i, Di=DNA quantity in fraction i, Ni=normalized abundance of the OTU per fraction i

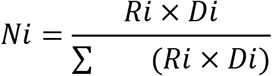

To detect enrichment, the weighted mean density of an OTU in the labeled and unlabeled samples was calculated and if the difference exceeded 0.005 g/ml the shift was determined significant (19). As quantitative SIP (qSIP) is sensitive to OTU abundance (19, 36), OTUs were used further only if they were non-spurious, whose distribution could not be differentiated from a normal distribution in both enriched and control samples (Kolmogorov-Smirnov test, alpha=0.05). With these criteria 180 OTUs were further analyzed.

### Metagenomic library preparation

The original unfractionated DNA extracted from the Sterivex filters was sheared by Covaris m2 to a mean length of 800 bp. Libraries were prepared from 15 ng of sheared DNA using the Ovation Ultra-low DR Multiplex System v2 kit (Nugen, Redwood City, CA, USA) with 9 amplification cycles. The libraries were bead purified as described above and sequenced on Illumina MiSeq for 600 cycles (UC Davis, USA) or on Illumina HiSeq rapid run for 500 cycles (USC genome core). See sup. table S1 for a detailed list of sequenced metagenomes.

### Metagenomic sequencing analysis

Reads were quality-trimmed using Trimmomatic version 0.33 as described above. Paired reads were assembled per sample in an iterative subsampling and assembly process as described in Hug et al. (37) but using metaSPAdes version 3.9.1 instead of IDBA-UD, followed by overlap assembly with minimus 2 with minimum overlap 200 bp and minimum identity 99%. Paired reads from all sequenced samples (sup. table S1) were mapped back to the contigs with BBmap (sourceforge.net/projects/bbmap/) requiring 95% identity. Binning was performed using two approaches: (1) binning the POLA 5/15 13C metagenome with CONCOCT (38) and bin refinement in Anvi’o (39). (2) pooling contigs longer than 5kbp from four naphthalene-enriched metagenomes (POLA 5/15 12C, POLA 5/15 13C, POLA 7/14 12C, POLA 7/14 13C), dereplicating them with cd-hit at 99% id (40, 41) and binning them using a combination of MaxBin2 (42), CONCOCT (38) and MetaBAT2 (43). These bins were combined, refined and reassembled using the MetaWRAP pipeline (44). Genomic bins generated by both methods were dereplicated using dRep (45) and only bins that were at least 50% complete and under 10% redundant were analyzed.

Initial taxonomic assignment of metagenomic assembled genomes (MAGs) was performed using GTDB-Tk (46). We then improved the taxonomy by generating class-level phylogenomic trees with GToTree (47) using NCBI RefSeq complete genomes and placing the bins assigned to the class by GTDB-Tk within them.

Reads from additional unenriched seawater metagenomes from all three sites were also mapped to the dereplicated MAG set to detect MAGs whose abundance increased in the presence of naphthalene.

### Metabolic analysis

Open reading frames (ORFs) in the final set of metagenomic assembled genomes (MAGs) and viral contigs were predicted using Prodigal (48) and annotated by assignment to clusters of orthologous groups (COGs) using the Anvi’o anvi-run-ncbi-cogs function. KEGG (Kyoto encyclopedia of genes and genomes) orthology for ORFs was assigned with Kofamscan using the prokaryote profile and its built-in thresholds (49). Kofamscan results were summarized using KEGGdecoder (50) (sup. fig. S2). ORF taxonomy was determined by Kaiju (51) using the RefSeq database.

Secondary metabolite clusters were identified using the antiSMASH bacterial version 5 online platform (52) with the “relaxed cutoffs” option. Iron-related transport and storage systems were identified using FeGenie (53). Additional functional analysis was also done with METABOLIC version 1.3 (54) and DRAM (55).

Read recruitment from different samples to the MAGs and viral contigs was analyzed with Anvi’o (39) using the Q2Q3 setting. This setting ignores the 25% lowest covered and 25% highest covered positions within the MAG when calculating mean coverage to avoid bias due to islands or highly conserved genes.

### Data availability

Metagenomic and amplicon raw reads from enrichment mesocosms can be found on EMBL-ENA under project PRJEB26952, samples ERS2512855-ERS2512864. The metagenomic library blank is under sample ERS2507713. Amplicon reads can be found under sample accession numbers ERS2507470-ERS2507679, PCR blanks and mock communities under samples ERS2507702-ERS2507712. Metagenomic t_0_ raw reads can be found under project PRJEB12234, samples ERS2512914-ERS2512919.

## Results

### Naphthalene uptake rates across the San Pedro Channel

Our main hypothesis was that at the Port of Los Angeles (POLA) there would be a seed of obligatory PAH-degrading bacteria due to chronic inputs. One indication to the existence of such a community would be measurable uptake of naphthalene-derived carbon. We also wanted to test whether this degradation potential extended out into the San Pedro Channel at the San Pedro Ocean Time-series (SPOT) and Two Harbors (CAT) (fig. 1). Indeed, isotopic enrichment measurements at POLA indicated a mean naphthalene uptake rate of 35 nM/d (standard deviation 19.53 nM/d) given a high input of 400nM naphthalene. However, naphthalene incorporation rates at SPOT and CAT were below detection.

### SIP-identified PAH-degrading taxa at POLA

^13^C-naphthalene-enrichment of POLA seawater led to significant incorporation of labeled carbon (>=0.005 g*ml^-1^ buoyant density increase corresponding to 9 atom %excess) by 34 out of 180 OTUs (sup. dataset S1). After 88 hours of incubation *Colwellia* sp. and *Cycloclasticus* sp. (Gammaproteobacteria) comprised 40% of the free-living (0.2-1 μ) microbial community. These OTUs were enriched at 53 (*Colwellia*) and 47 (*Cycloclasticus*) atom %excess (56) (fig. 2 A,B). These main naphthalene degraders were rare (<0.2% cumulative relative abundance of all OTUs classified as *Colwellia* or *Cycloclasticus*) to non-detectable prior to enrichment (t_0_) at all sites in all dates. While both taxa were represented by multiple OTUs, there was always a dominant OTU which matched the 16S-rRNA genes from the MAGs (see below). The most abundant OTU accounted for 95% and 99% of *Colwellia* and *Cycloclasticus* amplicons, respectively. Additional significantly enriched OTUs included Gammaproteobacteria (*Marinomonas*, *Neptuniibacter*, *Porticoccus, Pseudoalteromonas*, SAR86 and *Vibrio*), Flavobacteriales (*Tenacibaculum, Fluviicola, Polaribacter*, NS7 marine group and *Owenweeksia*), Sphingobacteriales, Deferribacterales (Marine group A), and Rhodospirillales (fig. 2 C-F).

**Figure 2:**
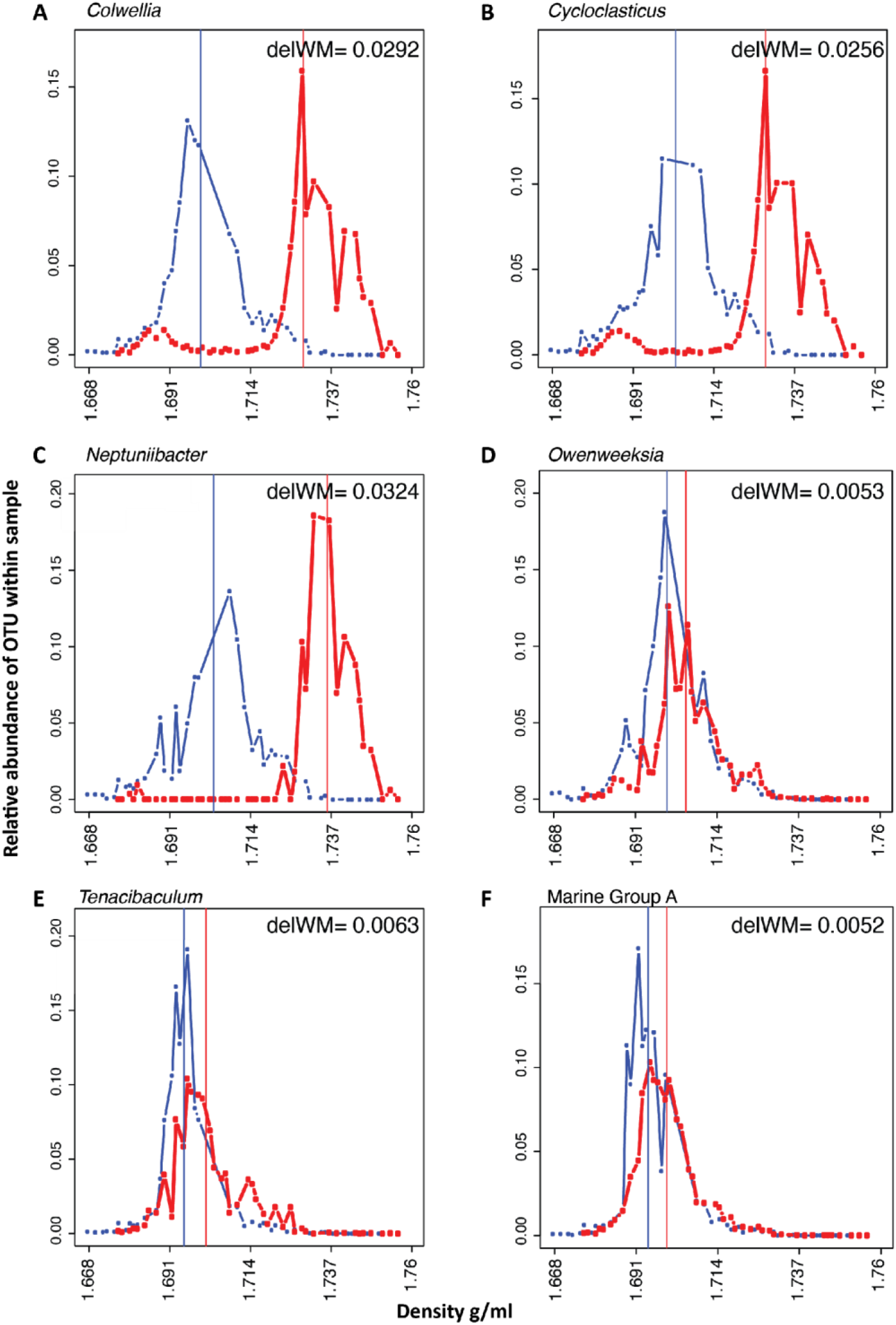
Density shifts demonstrating which taxa took up 13C naphthalene. Distribution of labeled (13C, red) and control (12C, blue) normalized relative abundance as a function of buoyant density of (A) *Colwellia*, (B) *Cycloclasticus*, (C) *Neptuniibacter*, (D) *Owenweeksia*, (E) *Tenacibaulum* and (F) Marine group A. Vertical lines represent the weighted mean of the distribution. The difference between the weighted mean density (WM) of the labeled (^13^C) and control (^12^C) (delWM) is noted on each plot.

### Metagenome-assembled genomes (MAGs) from naphthalene-enriched water

We assembled and binned 43 dereplicated MAGs that were more than 50% complete (mean=88%, SD=11%) and less than 10% redundant (mean=4.6%, SD=2.6%) from naphthalene-enriched POLA water. We then mapped reads from naphthalene-enriched and unenriched (t_0_) metagenomes to those bins in order to pinpoint potential degraders, under the assumption that potential degraders would be more abundant in mesocosm metagenomes compared to t_0_ metagenomes (fig. 3). Interestingly, seven bins were enriched in the presence of naphthalene at SPOT and CAT: POLA0515-13_bin_43, POLA0515-13_bin_23, Bin_4_4, Bin_47_1, Bin_46_1, Bin_20_1_1 and Bin_1_2_1 (sup. fig. S3). Eight bins significantly increased in abundance only in naphthalene-enriched POLA water (mean coverage < 1 in all CAT and SPOT samples) (table 1).

**Figure 3:**
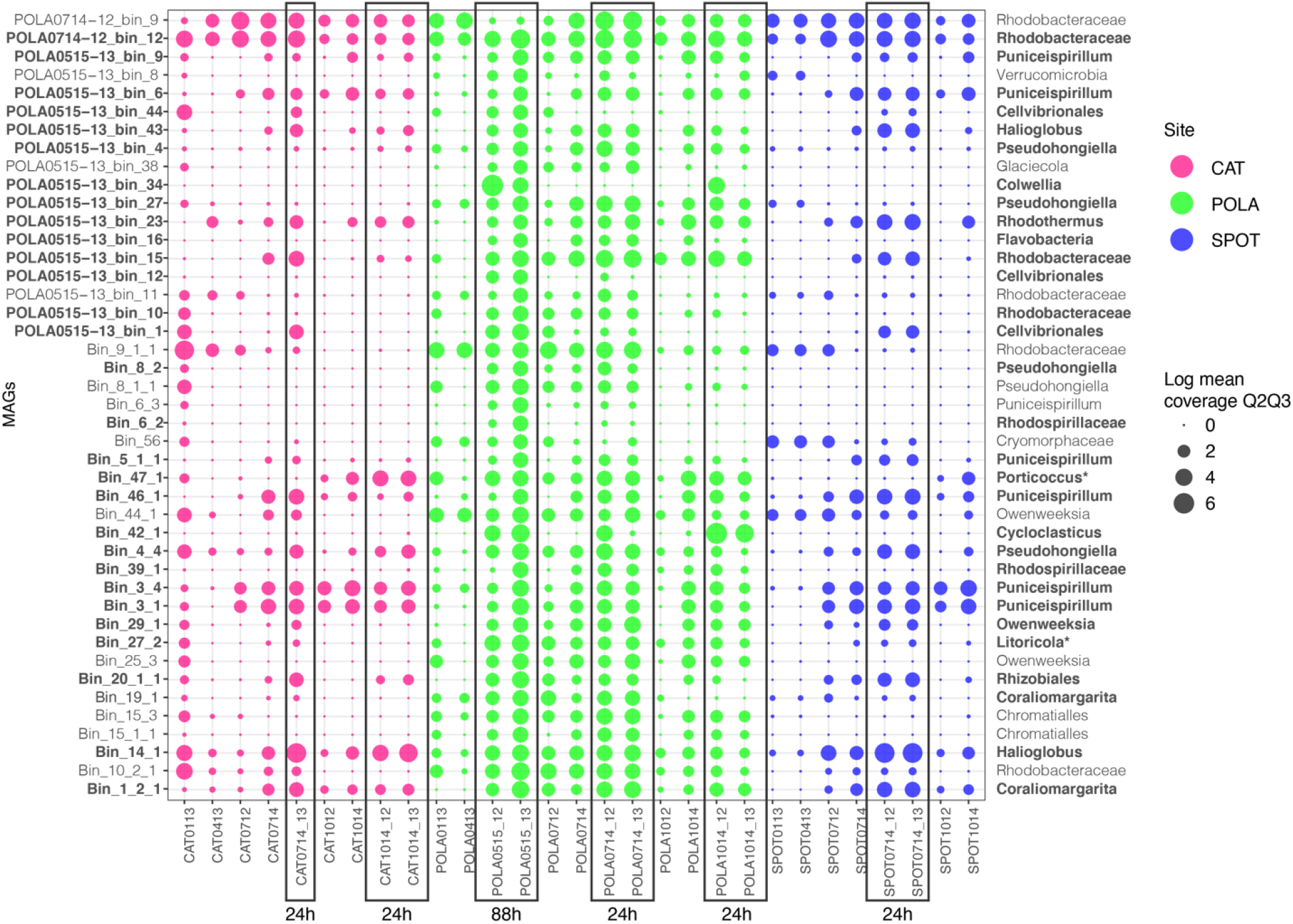
MAGs that respond positively to addition of naphthalene. Mean coverage of MAGs in naphthalene-enriched and unenriched seawater was normalized to sequencing depth and log-transformed. Each column represents a sample with collection month and year indicated (e.g. 0113 is January 2013). Enriched mesocosms have _12 (^12^C-naphthalene added) or _13 (^13^C-naphthalene added) next to the sample name and are surrounded by rectangles. Incubation time is denoted under the rectangles. The rest of the samples represent t_0_ (before naphthalene addition). MAG taxonomy by GToTree is displayed on the right. Bins in bold font had significantly higher coverage in naphthalene-enriched samples (p-value < 0.05). *16S-rRNA based taxonomy where its resolution was higher than GToTree taxonomy

**Table 1.** Taxonomy and genomic parameters of MAGs whose abundance increased at POLA upon naphthalene addition

| MAG ID | Taxonomy | Completeness | Redundancy | Size (Mbp) |
| --- | --- | --- | --- | --- |
| POLA0515-13_bin_34 | *Colwellia* | 91% | 3.6% | 3.5 |
| Bin_42_1 | *Cycloclasticus* | 99% | 2.9% | 2.4 |
| POLA0515-13_bin_16 | Flavobacteria | 99% | 3.6% | 2.7 |
| POLA0515-13_bin_12 | Cellvibrionales | 89% | 0.7% | 2.3 |
| POLA0515-13_bin_27 | *Pseudohongiela* | 93% | 10% | 2.8 |
| POLA0515-13_bin_4 | *Pseudohongiela* | 61% | 4.3% | 3.8 |
| Bin_39_1 | Rhodospirillaceae | 91% | 5% | 3.2 |
| *Bin_6_2* | Rhodospirillaceae | 96% | 4.3% | 3.8 |

### Primary naphthalene degraders

Two MAGs of interest were classified as *Colwellia* sp. and *Cycloclasticus* sp. and contained 16S-rRNA genes that matched at 100% identity to the most abundant OTUs of PAH-carbon incorporators (fig. 2 A,B).

The *Colwellia* MAG had high coverage and breadth (portion of the bin that has at least 1x coverage) only in naphthalene-enriched POLA water from May 2015 and October 2014 (fig. 3). Its closest relative based on a phylogenomic tree of 117 single-copy genes (47) is *Colwellia* sp. PAMC 20917 isolated from the Mid-Atlantic Ridge cold, oxic subseafloor aquifer (57) (sup. fig. S4; sup. dataset S2). It has a 78% average nucleotide identity (two-way ANI) and 63% average amino acid identity (two-way AAI) with the *Colwellia* MAG assembled from the Deepwater Horizon oil spill (18). This MAG contained subunits A and B of naphthalene dioxygenase enzyme (*nahAa*, *nahAb*), which is the first step in naphthalene degradation, but only parts of the remainder of the pathway known from pure cultures (e.g. *nahD, nahE*, salicylate hydroxylase large and small subunits, protocatechuate dioxygenase alpha and beta chains) (sup. dataset S3; S4). In addition, the MAG contained near-complete chemotaxis and flagella-assembly, near complete vitamin B6 and biotin biosynthesis, and a complete riboflavin biosynthesis pathway (sup. fig. S2). This organism has transporters for nitrite/nitrate, urea, phosphate, molybdate and heme. Finally, it can transport nitrite and potentially reduce it to ammonium via a dissimilatory pathway (*nirBD*, sup. dataset S3).

The *Cycloclasticus* MAG (99% complete; 2.9% redundant, 2.4 Mbp) was detected in POLA naphthalene-enriched metagenomes from May 2015, July 2014 and October 2014 (fig. 3 B). This MAG was most closely related to *Cycloclasticus zancles* 78-ME (sup. fig. S4; sup. dataset S2). It has a 78% average nucleotide identity (two-way ANI) and 81% average amino acid identity (two-way AAI) with the *Cycloclasticus* MAG assembled from the Deepwater Horizon oil spill (18). While it does not have the first steps of naphthalene degradation, it contains downstream genes of this pathway coding for 2-hydroxychromene-2-carboxylate isomerase (*nahD*), trans-o-hydroxybenzylidenepyruvate hydratase-aldolase (*nahE*) and catechol 2,3-dioxygenase (*xylE*) (sup. dataset S5). This MAG contains a near-complete flagellar assembly pathway but not the chemotaxis pathway. It can incorporate nitrogen from cyanate and biosynthesize riboflavin and biotin. This organism can degrade the aromatic hydrocarbon cymene and specifically contains the ring-opening enzyme *cmtC*. Similar to the *Colwellia* MAG, this MAG also contains transporters for nitrite/nitrate, urea and phosphate, and can potentially reduce nitrite to ammonium via *nirBD* (sup. dataset S3). The *Colwellia* MAG contains secondary metabolite clusters of homoserine lactone (hserlactone) and lassoprotein which imply quorum sensing and a potential antimicrobial activity respectively (58, 59). The *Cycloclasticus* MAG also contains clusters of bacteriocin and non-ribosomal peptide synthase (NRPS) which may also point to a potential antibacterial activity.

### Naphthalene-associated community metabolic characterization

In addition to *Colwellia* and *Cycloclasticus*, 27 MAGs were significantly more abundant in naphthalene-enriched samples (Wilcoxon rank test, p-value < 0.05). One of these MAGs, Bin_47_1 (*Porticoccus*) contained a 16S-rRNA gene that matched at 100% identity to an OTU that shifted significantly.

None of these MAGs contain the PAH or BTEX (benzene, toluene, ethylbenzene, xylene) degradation pathways defined by pure culture studies, but 12 of them had the KEGG pathway for degradation of the aromatic hydrocarbon cymene (sup. dataset S3). However, 18 MAGs included an aromatic ring-hydroxylating dioxygenase. Nine of these MAGs can assimilate nitrogen from nitroalkanes (fuel additives) or nitriles. These MAGs also contain secondary metabolite clusters of homoserine lactone (*Litoricola* Bin_27_2, Rhodobacteraceae POLA0714-12_bin_12), terpene (*Litoricola, Pseudohongiella*, Rhodobacteraceae, *Puniceispirillum*, SAR92, *Porticoccus*) and bacteriocin (Rhodobacteraceae). Finally, 24 out of these 29 MAGs contained at least one C1-oxidation enzyme, eight of the MAGs contain genes for oligosaccharide degradation such as alpha-L-rhamnosidase, beta-xylosidase and pullulanase and 23 MAGs contained tripartite ATP-independent periplasmic transporters (TRAP) (sup. dataset S3; sup. fig. S6).

Since several studies suggested that full degradation of PAHs may require bacterial consortia (18, 60), we also investigated the metabolic potential of the whole assembled community (open reading frames (ORFs) identified in contigs larger than 1kbp). The full naphthalene degradation pathway identified in pure cultures and incorporated into the KEGG database was not found even when combining all ORFs of the assembled MAGs.

Moreover, to confirm we did not miss additional naphthalene degradation genes because they did not assemble into contigs, we searched all forward reads from the POLA May 2015 ^13^C metagenome against all nucleotide sequences of naphthalene dehydrogenase ferredoxin subunit (*nahAc*, the first step of naphthalene degradation) from NCBI GenBank (188 references, September 2019). We found 248 reads (0.0008% of the total metagenome) that hit with a relaxed cutoff of e-value 10^-3^ (mean identity 97.8%, minimum bitscore 30). Using a conservative estimate that this gene comprises 0.1% of the metagenome (similar to 16S-rRNA), assuming one copy per cell and that each read maps to one copy of the gene, this would imply that only 8 out of 10,000 cells carry the naphthalene dioxygenase *nahAc* gene. This fraction would decrease even further if we dropped the single-copy and one read per gene assumptions.

A cross-comparison of the 29 MAGs that were significantly more abundant in naphthalene-enriched water showed that while most MAGs had the ability to synthesize hydrophobic amino acids (leucine, isoleucine, valine and proline), some polar amino acids (cysteine, histidine, serine, threonine) as well as charged amino acid lysine and amphipathic amino acid tryptophan, only one had the biosynthetic pathway for methionine, and none had biosynthetic pathways for arginine, phenylalanine or tyrosine. However, 15 MAGs had a putative transport system for polar amino acids or branched amino acids, and nine had a transport system for L-amino acids. Only five MAGs contained a urea transport system, and while 34% naphthalene-enriched genomes had a urease gene, so did 43% of the genomes that did not respond to naphthalene-enrichment. In addition to *Colwellia* and *Cycloclasticus*, eight MAGs can degrade urea to ammonia via urease and four MAGs have genes coding for *nirBD* nitrite reductase. Five MAGs contain a phosphonate transport system whereas 20, including *Colwellia* and *Cycloclasticus*, can transfer phosphate.

Regarding vitamins, in addition to *Colwellia*, *Cycloclasticus*: four MAGs can synthesize biotin and 12 MAGs can synthesize pantothenate (vitamin B5). *Colwellia* and four additional MAGs can synthesize riboflavin (vitamin B2) (sup. fig. S2, sup. dataset S3).

Of these 29 genomes five are capable of anoxygenic photosynthesis (three Rhodobacteraceae and two *Halioglobus}*, and 18 encode proteorhodopsins (sup. dataset S3).

## Discussion

### Naphthalene biodegradation potential does not extend outside the Port of Los Angeles

Naphthalene degradation rates were detectable only at POLA but not at SPOT or CAT. As there is tidal mixing across the San Pedro Channel, we would expect some potential for degradation at those sites by bacteria advected from the port. A possible explanation for the non-detectable rates at SPOT and CAT is that diminishing PAH inputs offshore (61) can support a very limited degrader seed community. The PAH degraders would then be out-competed for nutrients at those sites by microbes that are better adapted to oligotrophic conditions, and lose viability or be otherwise removed by processes like grazing and viral infection at rates faster than they are replenished by mixing or growth. Indeed, the abundance of MAGs of the primary degraders *Colwellia* and *Cycloclasticus* was extremely low at CAT and SPOT. More studies quantifying naphthalene degradation rates across San Pedro Channel are needed to determine whether this pattern consists throughout the year. The PAH carbon incorporation rate at POLA, however, indicated a minimum removal of ~10% of the initial concentration per day. This rate is roughly 3-fold higher than that measured in seawater from the Gulf of Mexico amended with crude oil (6, 62), although the incubation time was shorter compared to the Gulf of Mexico experiments, and the degradation rate may not be linear. The rate we measured is likely underestimated as measurement was performed using GF/F filters, which have a pore size leading to loss of roughly half of marine free-living prokaryotes. In addition, there could be degradation of PAH without incorporation of the labeled carbon into cells, which would not be accounted for by this measuring technique.

### Naphthalene-associated community

Incubation of POLA water with naphthalene revealed a difference in PAH associated communities between the 24-hour and 88-hour incubations. First, it is noteworthy that many of the MAGs that were abundant in naphthalene-enriched cosms after 88 hours were already abundant after 24 hours. Some of them were common marine heterotrophs such as *Puniceispirillum* (SAR116) and Rhodobacteraceae. Both *Pseudohongiella* and OM182 MAGs were matched by 16S-rRNA gene to OTUs that did not incorporate carbon from naphthalene, indicating that their increased abundance in the presence of naphthalene is not due to them being primary degraders. We speculate that their increased abundance in enriched cosms is due to an early response to the presence of naphthalene, as their initial abundance was higher than that of the primary degraders *Colwellia* and *Cycloclasticus*. However, we also note a difference in response time between the primary degraders. In 3 out of 4 24 hours incubations, *Cycloclasticus* abundance was already comparable to its abundance after 88 hours, whereas *Colwellia* was only abundant in one 24 hours incubation, and its abundance was orders of magnitude lower compared to 88 hours. This may indicate a difference in growth rates and/or utilization of substrates and nutrients, as both taxa were undetectable in unenriched water. Guttierez et al. (6) identified 4-5 orders of magnitude increase in abundance of *Colwellia* and *Cycloclasticus* in PAH enriched mesocosms after three days. It is notable that the same naphthalene degrading taxa appeared reproducibly in multiple incubations in different seasons. Thus, characterization of the metabolic requirements of these specific MAGs could be helpful in the future in case of an oil spill at the Port of LA.

Second, some MAGs were abundant in enriched water from SPOT and/or CAT even though there was no measurable degradation of naphthalene. They could be benefitting from toxicity of naphthalene to other bacteria or from byproducts of naphthalene oxidation. In the case of byproducts utilization we would have expected OTUs associated with these MAGs to shift to a heavier density. Only one of these MAGs representing *Porticoccus* contained a 16S-rRNA gene, and the matching OTU indeed shifted.

Incubation conditions were not identical between the 24-hour cosms from all three sites in July and October 2014 and the 88-hour cosm (POLA only, May 2015). Incubation in the dark (88 hours) would promote heterotrophy while incubation in the light should still allow cyanobacteria and photosynthetic picoeukaryotes to compete over nutrients with potential naphthalene-associated bacteria. However, if light had affected the results, we would have expected it to be reflected in MAG abundance. As it is not, we suggest that under naphthalene enrichment these photoautotrophs are not good competitors.

The duration of the 88-hour SIP experiment may have contributed to the accumulation of labeled carbon in bacterial DNA. In order to observe pronounced enrichment, the PAH degraders have to replicate at least once, and DNA density increases with every replication as the new strand is made with labeled nucleotides. Additionally, it is methodologically difficult to observe enrichment in rare taxa (36). Since the seed degrader community is made up of rare organisms, their abundance must increase substantially before their enrichment can be tracked. The other side of the coin is that a longer incubation increases the chance of cross-feeding, namely labeling of organisms that incorporate intermediate- or end-products of naphthalene degradation. Crossfeeding of intermediate products of naphthalene degradation is a vital part of the complete remineralization of this hydrocarbon which appears to have been often overlooked. In this study, there were 24 OTUs which did not belong to the main naphthalene degraders (*Colwellia*, *Cycloclasticus* and *Neptuniibacter*) but were still significantly enriched (9-15%). While two of them (i.e. *Marinobacter* and *Pseudoalteromonas*) may be involved directly in degradation of naphthalene but either grow more slowly or were simply outcompeted by *Colwellia* and *Cycloclasticus* (63, 64), the rest are more generalist heterotrophs which are also associated with degradation of algal blooms in the San Pedro Channel (e.g. Flavobacteria, Marine group A, SAR86) (65).

### Metabolism of putative PAH-degraders and associated taxa

In order to gain more insight into metabolic requirements of naphthalene degrading bacteria at POLA, we examined metabolic pathways and key transporters and enzymes within the annotated proteins in our metagenomic assembled genomes (MAGs). To date, only one published study characterized metabolic pathways in assembled genomes of marine oil degrading bacteria, using metagenomes from naphthalene- and phenanthrene-enriched seawater from the Deepwater Horizon (DWH) oil spill (6, 18). Within the DWH mesocosms, the prominent naphthalene degraders belonged to the genera *Cycloclasticus* and *Alteromonas*, and phenanthrene degraders included *Neptunomonas*, *Cycloclasticus* and *Colwellia*, whereas we found *Cycloclasticus* and *Colwellia* to be the primary naphthalene degraders at POLA.

Both *Colwellia* and *Cycloclasticus* were non-detectable in both 16S-rRNA amplicons and MAG coverage before enrichment. However, after 88 hours of incubation they were the most dominant taxa in the mesocosms, and exhibited very significant enrichment indicating incorporation of naphthalene-derived carbon into their DNA over a substantial number of replication cycles. The *Colwellia* MAG was the only one assembled here that contained annotated subunits of naphthalene 1,2-dioxygenase (*nahAa*, *nahAb*). As this is the first step of the pathway, requiring investment of reducing power (66), it is not surprising that some of the downstream metacleavage of catechol is also present in the MAG. In comparison to the naphthalene degradation enzymes found in the *Colwellia* genome from the Gulf of Mexico, this MAG also had subunit A of naphthalene dioxygenase (*nahAa*) as well as *nahD*. It had an additional subunit (*nahAb*) of naphthalene dioxygenase but lacked the next enzyme encoded by *nahB*.

*Cycloclasticus* genes dominated the metagenome of the phenanthrene-enriched DWH mesocosm (18), highlighting its importance as an obligate PAH degrader (67). Unlike the *Colwellia* MAG, our *Cycloclasticus* MAG contained hardly any part of the KEGG-defined aerobic naphthalene or phenanthreme degradation pathways which are based on pure cultures. This is in stark contrast to the *Cycloclasticus* genome assembled from naphthalene-enriched water from the Gulf of Mexico (18). However, it did include several ring-hydroxylating dioxygenases. While these enzymes are best known to degrade single aromatic hydrocarbons, previous studies demonstrated that some single-ring aromatic degrading enzymes are capable of degrading naphthalene (two aromatic rings) efficiently (67–70). Additionally, the *Cycloclasticus* MAG has a sigma-54-dependent transcription regulator with a potential hydrocarbon-binding domain as identified in *Cycloclasticus zancles* 78-ME (GenBank accession AGS40441.1), which could control transcription of hydrocarbon degradation genes. Dioxygenases require iron and an iron-binding domain, such as ferredoxin which can be shared by multiple enzymes (71, 72), which we found in this MAG. As we know that our *Cycloclasticus* strain can take up naphthalene-derived carbon as its main carbon source based on the SIP results, and has genes that can participate in similar pathways but none of the culture-defined pathway, we posit that non-traditional dioxygenases with affinity to naphthalene were utilized to degrade naphthalene. It is reasonable to expect that generalist aromatics-degrading organisms might “most-economically” possess multi-functional pathways that do not fully optimize degradation of a single substrate but work adequately with several aromatic substrates, without the need to carry the “extra baggage” needed to optimize each specific pathway. To further support this idea, metatranscriptomes of the microbial community within the oil plume of the Deepwater Horizon spill revealed low to nonexistent transcription of known PAH-degradation genes despite their presence in metagenomes (73). Moreover, the use of stable isotope probing indicated that both *Colwellia* and *Cycloclasticus* had the ability to degrade PAHs, whereas if only metagenomics was used, we might have concluded that they did not.

The presence of secondary metabolite clusters in both genomes suggests that part of the success of these organisms may be attributed to antimicrobial activity in addition to the ability to incorporate carbon from naphthalene.

Interestingly, most enriched MAGs also had Tripartite ATP-independent periplasmic (TRAP) transporters. TRAP transporters are common in marine bacteria (74) have been shown to participate in chlorobenzoate (75) and lignin-derived aromatic degradation (76), suggesting that they may participate in uptake of PAHs or their byproducts in this system. The presence of C1-oxidizing enzymes in most naphthalene-enriched MAGS also implies that even those who cannot directly degrade naphthalene may benefit greatly from degradation byproducts.

### MAG-generated hypotheses for future biostimulation experiments at POLA

Previous studies revealed that in many systems bioremediation of crude oil can be enhanced by addition of nitrogen and phosphorous (15, 77). While oil degrading bacteria are found in many marine systems (78), their metabolic requirements may vary by system, depending on limitations *in-situ*. The significant genomic dissimilarity between the strains of *Colwellia* and *Cycloclasticus* found at POLA and the ones found at the Gulf of Mexico implies that we cannot assume similarity in metabolic requirements between systems. As oil degradation requires a considerable amount of nitrogen (79), choosing the correct substrate could be crucial.

Both *Colwellia* and *Cycloclasticus* displayed a potential for using a variety of nitrogen sources to different ends, with a full dissimilatory nitrite reduction to ammonium (DNRA) pathway and nitrite/nitrate transporters (*focA*, *narK*) as well as a urea transporter and the ammonium-assimilating glutamate synthase pathway. Nitrate, nitrite and ammonium are always detectable at POLA in surface seawater (sup. fig. S5), and thus are not limiting nutrients in this site. DNRA is an anaerobic process, which appears to be in conflict with the aerobic nature of surface seawater. However, PAHs are hydrophobic and tend to attach to particles, and particles can provide anaerobic microniches in their interior (80). It is possible that PAH degradation occurs in large part on particles at POLA, which would explain the presence of this pathway within the MAGs, and that in these taxa nitrite is sometimes used as an oxidizer whereas ammonia and/or urea are used as nitrogen sources. *Colwellia* and *Cycloclasticus* strains have also been isolated from sediments, where conditions may become anaerobic and support DNRA (8, 10).

Based on the functional pathways and ABC transporters found in MAGs of naphthalene incorporators, we propose specific targets for future experiments on enhancement of PAH bioremediation at the Port of Los Angeles. While it appears both local strains of *Colwellia* and *Cycloclasticus* can utilize urea, when considering the remaining MAGs enriched in the presence of naphthalene, addition of ammonia rather than urea may be more beneficial in our system. Similarly, phosphate is more likely to augment bioremediation than phosphonate. To supply iron for dioxygenase synthesis, adding heme/hemoproteins should be superior to addition of Fe(II) or Fe(III) as 27/29 enriched MAGs contain heme transporters. Marine bacterioplankton have been shown to be able to incorporate iron from heme groups (81, 82). Finally, Polar amino acids for which we did not find any biosynthetic pathways but did find transporters in enriched MAGs could also potentially augment bioremediation.

In the current climate of excessive use of fossil fuels, chronic deposition of toxic and recalcitrant polycyclic aromatic hydrocarbons into the coastal ocean is inevitable. PAH-degrading bacteria may provide some control over the remineralization of these inputs and could serve as targets for bioremediation technologies. Identification of naturally-occurring biodegraders is a crucial first step, but optimization of the degradation process requires knowledge of the metabolic requirements of local organisms (23). Moreover, their genomic information remains an available resource should other hypotheses for biostimulation arise. The combination of stable isotope probing with metagenomics could provide a source for genomically-generated hypotheses and experimental design for future bioremediation experiments.

## Acknowledgements

We extend our gratitude to Prof. Douglas Capone (USC) and Troy Eric Gunderson for the uptake rate measurements and for comments on this manuscript. We would like to thank Dr. Ben Tully for assistance with analysis of functional pathways. This research was made possible with the support of NSF grants 1136818 and 1737409, Gordon and Betty Moore Foundation Marine Microbiology Initiative grant GBMF3779 and The Wrigley Institute for Environmental Studies Norma and Jerol Sonosky fellowships to E.T.S.

## Competing interests

The authors declare no competing interests.

## Supplementary figures titles

Figure S1: Relative photosynthetically active radiation (PAR) profiles at the port of Los Angeles (POLA), the San Pedro Ocean Time-series (SPOT) and Santa Catalina Island (CAT) in spring (April), winter (January) and fall (October)

Figure S2: Basic metabolic pathways in MAGs represented as fraction of the enzymes present. *Colwellia* and *Cycloclasticus* MAGs are labeled with a star.

Figure S3: Mean coverage of MAGs with increased abundance in naphthalene-enriched water from SPOT/CAT

Figure S4: Phylogenomic tree showing the placement of our *Colwellia* and *Cycloclasticus* MAGs. The MAGs are labeled in purple. The tree is based on 117 single-copy genes of Gammaproteobacteria and was built in GToTree. Complete reference genomes were downloaded from RefSeq on 8/20/2019.

## Supplementary table titles

Table S1: Sample details – number of metagenomics (MG) reads and 16S-rRNA amplicons. * Number of amplicons in enriched mesocosms are combined across all density fractions.

## Supplementary dataset titles

Dataset S1: p-values of Kolmogorov-Smirnov normality test of ^12^C- and ^13^C-naphthalene enriched OTU distribution as a function of density, density shift, corresponding atom% excess and SILVA-assigned taxonomy of 180 OTUs analyzed from POLA 5/15.

Dataset S2: General information on metagenomic assembled genomes (% completion, % redundancy, length), taxonomy determined by GTDB-Tk, Anvi’o, GToTree and 16S-rRNA and mean coverage in metagenomes (second and third quartile – Anvi’o setting Q2Q3)

Dataset S3: Results of METABOLIC analysis of the 29 MAGs whose abundance increased in the presence of naphthalene. Each column represents a MAG, and each row represent a Kegg module.

Dataset S4: Results of DRAM hits to open reading frames of the *Colwellia* MAG and *Cycloclasticus* MAG

Dataset S5: Results of DRAM hits to TRAP systems of the 29 MAGs whose abundance increased in the presence of naphthalene.

